# The morning burst: Shifting daily patterns of ATP production in Drosophila and temporal windows for their improvement in ageing

**DOI:** 10.1101/2021.09.14.460393

**Authors:** Harpreet Shinhmar, Jaimie Hoh Kam, John Mitrofanis, Chris Hogg, Glen Jeffery

## Abstract

Mitochondria produce energy for cell function via adenosine triphosphate (ATP) and are regulated by a molecular 24h clock. Here we use *Drosophila melanogaster* to reveal shifts in whole animal ATP production over 24h, showing a marked peak in the morning that declines around midday and remains low from then through to the following morning. Mitochondrial membrane potential and ATP production has been shown previously to improve after long wavelength exposure, but apparently not at all times. Hence, to explore this further we exposed flies to 670nm at different times. Exposures between 08.00 and 11.00 resulted in a significant increase in ATP, while exposures at other times had no effect. Within the morning window, not all times were equally effective, however, 670nm exposure mid-morning when ATP production was maximal did not increase ATP, possibly because mitochondria lacked spare capacity at this time. Hence, in the morning there is a complex dynamic relationship between long wavelength light and mitochondria. Mitochondrial function and the influence of long wavelengths are conserved across species from fly to human, and determining the time points for light administration to improve function in ageing and disease is of key importance. Our data progress this search and reveal the outline of these times.

## Introduction

Many physiological processes are regulated over a 24-hour period to adapt to environmental changes of day and night (1,2). Such regulation includes the sleep-wake cycle, locomotor activity and feeding during the active phase (3–11). It is able to keep running even under constant 12 /12-hour light/dark environmental conditions.

There is evidence that the function of mitochondria, the major metabolic hubs that produce ATP for cellular function, is also regulated by a 24-hour clock (12) This is not only in terms of their metabolic function but also in terms of their complex dynamics that, include fusion and fission processes that are central to the maintenance of their integrity and whose disruption is associated with ageing and disease (13–15).

With age and many diseases mitochondrial respiration declines (16) and their circadian behaviour is disrupted (13). However, aged mitochondrial respiration can be improved with longer wavelengths of light (650-900nm). This range of wavelengths has been shown to improve mitochondrial membrane potential (17) and ATP production (18). These improvements have been associated with increased longevity and neuroprotection in ageing in insects and mammals (19–21). Recently, it has been reported that such improvements cannot be achieved consistently over the 24-hour period, and that there may be an optimum period of exposure that selectively improves mitochondrial function (22). The precise time point(s) during the day for generating the maximum impact for improving mitochondrial function is not known. In this study, we determine the shifting pattern of mitochondrial activity by investigating ATP production over a 24-hour period in whole aged flies. Thereafter, we explored whether there are specific periods of sensitivity when long wavelength light (670nm) is most effective in improving mitochondrial function and periods when it is ineffective. Our results are key to better understanding of overall mitochondrial function and when this function can be improved in ageing and potentially disease using long wavelength light.

## Materials and methods

### Fly stocks and husbandry

The wild-type, male *Drosophila melanogaster* Dahomey flies were used throughout this study. Newly hatched male flies were collected and kept at a density of 30 flies per food vial. They were maintained under standard laboratory conditions and diet with 12/12 light dark (LD) cycle at 25° C and 70% humidity. Five week old flies were used throughout the study. This age was selected because older flies suffer mitochondrial decline and hence are suitable for experiments where mitochondrial function can be improved with long wavelength light. All flies were culled by instant freezing with dry ice.

### ATP

ATP measurements were undertaken using a commercially available ATP determination assay (ThermoFisher Scientific, UK). Flies were collected at 30min intervals between 03.00 - 21.00 and at 1h intervals between 22.00 – 03.00. There were 5 flies per replicate, with a total of 7 replicates per time point were homogenized in extraction buffer consisting of 6M guanidine-HCL in 100mM Tris and 4mM EDTA, pH 7.8, followed by a heat treatment at 95 0C for 5 min. The homogenates were then centrifuged at 16,000 X g for 15 minutes at 4 0C and the supernatant was collected. The protein concentration of each sample was measured using the BCA assay (ThermoFisher Scientific, UK). For ATP measurement, samples were diluted in 1/10 with the extraction buffer and were added to the reaction mix prepared according to the manufacturer’s instructions.

### 670nm light treatment

Flies were exposed with 670nm LED light devices (C.H. Electronics, UK) for 15 minutes daily for 7 days at 40mW cm^-2^ (at 08.00, 09.00, 10.30, 11.00, 16.00 and 05.30). Each experimental group had a matched control whereby they were exposed to the same 12/12-hour lighting. The fly vials were examined with a spectrometer and a radiometer prior to 670nm exposure. They reduced the overall energy applied by <5% and did not modify the wavelength.

### ADP/ATP

Flies exposed to 670nm were processed for changes in both ATP and the ADP/ATP ratios. For ADP measurement, ATP was measured, then an ADP-Converting enzyme (Abcam, UK), which converts ADP to ATP was added according to the manufacturer’s instructions. The ADP levels are calculated by subtracting the second reading from the first.

## Results

### Shifts in ATP production during 24-hour cycle

ATP levels in 5-week-old flies were assessed every 30 minutes from 03.00 to 22.00 and at hourly intervals during the other period (Figure 1). The results show that as the light came on at 06:30, ATP level were rising, peaking at around 10:30. Thereafter, levels declined slowly and by 14.00 had started to plateau over the course of the day. When the lights were turned off at 18:30, there was a further decrease in ATP production, before the levels plateaued finally at around 03:00 until the light turned on again at 06:30.

**Figure 1.**
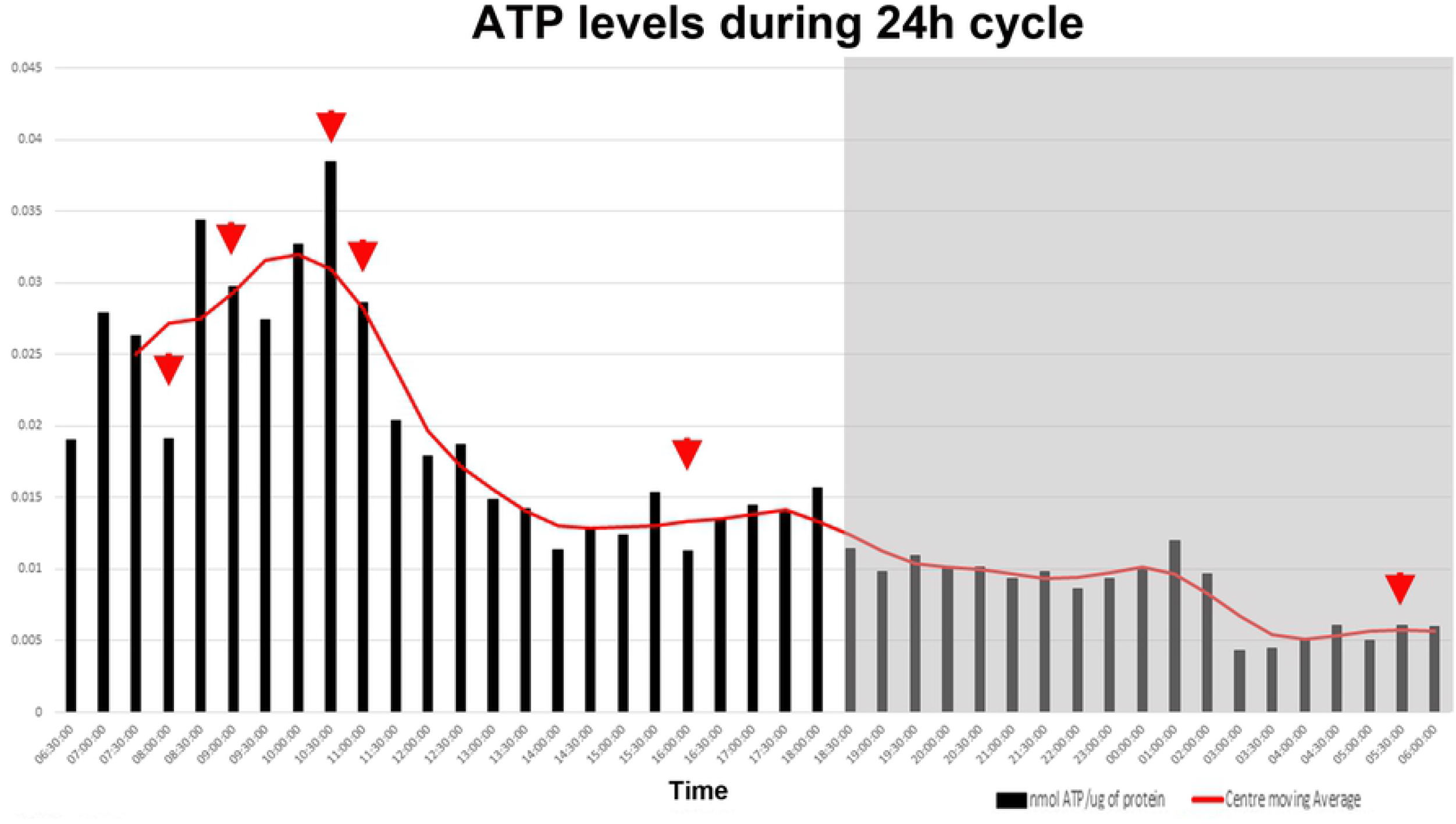
Shifting ATP levels during 24-hour cycle in 5-week-old flies. ATP levels were measured in flies every half an hour during light ON (daytime) and every hour during light OFF (night-time). The graph shows that as the light is ON at 06:30 the ATP level starts rising to peak at around 10:30 then slowly slopes down after. There is a further decrease in the level of ATP when the light is OFF at 18:30 to trough at around 03:00 and plateau until the light returns on at 06:30. Red arrows are selected time points where the flies were treated with 670nm light. Light OFF phase is shaded in grey.

### Effect of 670nm red light on ATP levels during 24-hour cycle

In order to examine the effect of 670nm light on the varying patterns of ATP production, flies were treated for 15 minutes daily over 7 days (Figure 2) at six different time points that corresponded to periods of change in the overall 24h patterns of ATP production (Figure 1). The first two time points were at 08.00 and 09.00, on the ascending slope of the peak of ATP production in the morning. Third was on the top of the slope when the ATP level was at its maximum at 10.30. The fourth was at the descending slope in the morning at 11:00. The fifth, corresponded to the early phase of the afternoon plateau at 16:00 and sixth in the dark phase at 05:30 when ATP production was close to its minimum.

**Figure 2.**
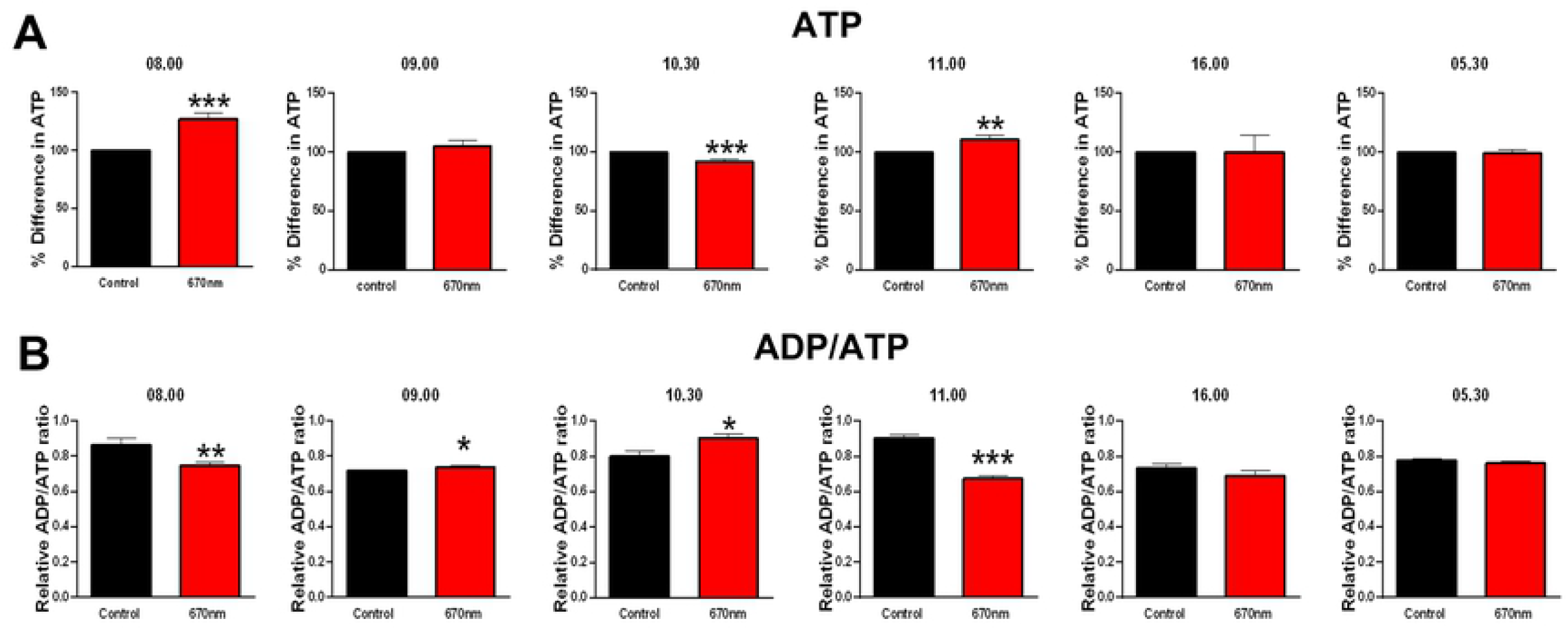
**A**. 5 week old flies were then treated at different time points with a 670nm light for 15 minutes daily over 7 days, and the level of ATP was measured and compared to their age-matched controls. There were no significant changes in the level of ATP at 09:00 (↑ 5%), 16:00 (↓ 0.3%) and at 05:30 (↓ 0.7%), but there were significant increases in the ATP level when the flies were treated at 08:00 (P = 0.0003, ↑27%) and 11.00 (P=0.0084, ↑11%) corresponding to the ascending and decreasing slope respectively. There was a significant reduction in the ATP level at 10.30 (P=0.0003, ↓8%) when the ATP was at its natural optimum level (Figure 1). There were no changes in ATP level when flies were treated at 16.00 and 05.30. **B.** The ADP/ATP ratios were also measured to provide insight into the overall ATP pool during the times of red light exposure. There was a significant increase in the ADP/ATP ratio in the flies treated with 670nm light at 09:00 (P= 0.0265) and 10.30 (P=0.0189) compared to the controls. There was a significant decrease in the ratio of ADP/ATP levels in flies treated at 08.00 (0.0051) and 11:00 (P=0.0003). There were no change in the ADP/ATP ratio when flies were treated at 16:00 and 05.30. Abbreviations: * p<0.05,, ** p<0.01, *** p<0.001. ↑ = increase, ↓ = decrease.

Changes in ATP production as a consequence of 670nm light exposure were confined to the morning period. However, even within this period, 670nm applied at different exposure times did not result in consistent changes, indicating that a more complex relationship exists between 670nm light exposure and mitochondrial function than simply a sensitive/insensitive switch. There were significant increases in ATP at 08.00 (P = 0.0003, increase of 27%) and 11:00 (P = 0.0084, increase of 11%). However, at the peak of ATP production at 10.30 (Figure 1), 670nm light exposure was associated with a decrease in ATP levels. This may be because at this point, mitochondrial ATP production was maximal and there was no spare capacity for its further elevation. There were no significant changes in the level of ATP after 670nm exposure at either 09:00 (increase of 5%), 16:00 (decrease of 0.3%) or 05.30 (decrease of 0.7%) (Figure 2A). While the exposure at 09:00 did not result in a significant increase in ATP, the 5% change at this time point was greater than found at other periods where ATP could not be elevated by 670nm light exposure. It is possible that 09:00 is a transition period between relatively effective and ineffective periods of sensitivity.

### Shifting adenosine diphosphate (ADP) levels during 24-hour cycle and the effect of 670nm light exposure

Finally, we examined whether the changes in ATP production during the day were mirrored also by changes in ADP by examining ADP/ATP ratios at times when 670nm was applied. Overall, our results showed that if ATP was not changed by 670nm exposure, neither was the ADP/ATP ratio. However, when ATP was elevated during the 24hr cycle, the ADP/ATP ratio declined and when ATP was reduced, the ADP/ATP ratio increased. The flies treated at 08.00 and 11:00 showed a significant decrease (P= 0.0051 and P= 0.0003 respectively) in ADP/ATP ratios, likely to be due to ADP conversion to ATP, and a release and utilisation of cellular energy.

There was a small, but significant increase in ADP/ATP ratios in flies treated with 670nm red light at 09:00 (P = 0.0265), reinforcing the notion that this time point may be one of transition and a significant increase at 10.30 (P = 0.0189) compared to controls (Figure 2B). Hence, at these time points it is likely that ATP was being converted to ADP, potentially as an energy storage mechanism. There were no significant differences in ADP/ATP ratios at the other times examined, 16.00 and 5.30 (Figure 2B) when 670nm light had no impact on ATP levels (Figure 2A).

## Discussion

This study had two major findings. First, that total ATP levels in flies vary considerably over a 24h period, peaking in the early part of the day associated with onset of light. Second, during this morning peak period, there were particular time points where it was possible to manipulate ATP levels optimally with 670nm light exposure. However, the morning peak period contained a number of complexities in addition to ATP levels undergoing changes over time, as there were changes in the sensitivity of mitochondrial responses to 670nm exposure as well, at least in terms of ATP and ADP/ATP ratios. Hence, a key limitation of this study is that it examines a highly dynamic situation with fixed time point analysis. However, our findings provide key data for where future analysis should be focused.

Several previous studies have reported that mitochondrial biogenesis (23,24), content (25,26), dynamics (27), and function (28–30) are regulated by over a 24-hour period. These studies however, were focussed on single organs, for example liver, brain and muscles or cultured cells, indicating that the mitochondria within each organ set their own autonomous clocks (31). Hence, our study differs from many previous investigations by examining the whole animal. But even within single organs there are variations. Yamazaki et al. (32) measured ATP levels in 3 regions of the rat brain over 24-hour and showed variations in peaks of production. Although the key difference was in the superchiasmatic nucleus compared to the two other regions, both in timing and in the levels of ATP production, the authors suggested that that variation was due to the different activity patterns and energy demands of each region. In essence, targeting one organ does not give one an overall understanding of how ATP levels fluctuate over a 24-hour period in a whole organism and this is why we chose to use the *Drosophila* fly model.

Our results showed that flies elevate ATP production early in the day, peaking around 10:30. This may be for the normal active physiological function for the rest of the day. Hence, flies increase mitochondrial function when lights were turned on, presumably related to an increased energy demand and resumption of normal activity. This result may be reflective of their overall feeding patterns. Xu et al. 2008 (33) have reported that, flies start to feed prior to or around the time of lights ON in the morning but then decrease their feeding behaviour between 4-6-hours after lights ON. This matched closely our peak ATP level at 10:30, which is 4.5-hours after lights ON and the general decline in levels thereafter. Further, Xu et al. 2008 (33) showed a small peak in the fly feeding pattern at 8-10-hours after lights ON just before onset of night, which is reflected broadly here in our finding of a small peak at 17:30.

Dubowy et al. (34) traced *Drosophila* fly locomotor activity over 24-hour and have shown two activity peaks, one corresponding to dawn (at lights ON) and the second at dusk (at lights OFF), with the flies anticipating these events and increasing their activity just before day-time and night-time. This may explain our finding of a dip in ATP levels between 03:30 to 06:30, as the flies at these times may have a high ATP consumption rate. This patterns aligns with our previous findings showing an increase in ADP levels and glycolysis during this precise time period in flies (35). Our results also showed ATP levels peak at 10:30, during the period when locomotor activity was measured at its lowest point (30), which may be explained by the flies having a low ATP demand at this point. The second peak in locomotor activity recorded just before night (30) appears associated with an additional decrease in ATP shown her; hence unlike the morning burst, the flies seem to be utilising the energy available in the afternoon.

Finally, our results showed that exposure to 670nm light improves mitochondria function effectively, but only within a limited time window, between 08:00 and 11:00. Importantly 670nm is completely ineffective at time points sampled outside this time window. Weinrich et al. (35) examined a number of mitochondrial metrics over a limited series of time points over the day in flies, and revealed shifts in mitochondrial complex activity, glycolysis and animal respiration. They also showed that 670nm light exposure increased respiration rate when given in the morning but not the afternoon. When the ATP levels of aged flies were improved by a morning 670nm light exposure it translated to increased mobility, cognition and retinal function (18). Similar patterns have been found in aged human retinal function, with morning 670nm light exposure improving colour contrast ability that was not possible when exposures were in the afternoon (Shinhmar *et al*. Submitted). Hence, these mechanisms in mitochondria are likely to be conserved (31). This highlights the need to define the temporal windows in which ATP can be improved in aged or damaged systems, given the interest in the use of long wavelength light and its potential. The results presented here take a step in refining the answer to this problem.

There are two key implications for the results that we present. First, mitochondria decline is a key feature not only in ageing and disease, but also in insecticide poisoning that undermines pollinators that play a critical role in agribusiness. Previously we have shown that bees poisoned with nicotinoid insecticides are protected by exposure to 670nm light by improving mitochondrial function, immunity respiration and visual function (36,37). If protection is to be maximised, particularly in the translations from the lab to the more complex environment of the field, understanding the temporal windows for long wavelength light application is of major importance. In this respect very similar patterns of effective light exposure are being revealed when lights are placed in hives in the field by Beefutures (France). Honey bees in these hives show improved resistance to environmental challenge that undermines mitochondrial function.

The therapeutic implications of our results are considerable. Red to near infrared light has been shown by many studies to improve cell function and survival in animal models of ageing (19) (38–40) and of a range of neurological conditions, for example macular degeneration, Parkinson’s disease, Alzheimer’s disease, traumatic brain injury, stroke and multiple sclerosis (41–48). There are also clinical reports of positive outcomes in aged humans and in patients suffering from many of these conditions (49–56). Our findings, coupled with the highly conserved nature of the mechanism across species indicated that there are specific but limited times of day when this method is effective, but long periods when it is not; this feature could prove pivotal in achieving the maximum therapeutic outcome in both experimental laboratory and clinical settings.

## Funding body

This work was supported by Beefutures, France.

